# Terrestrial ecosystems enhance root zones in response to intensified drought

**DOI:** 10.1101/2024.05.26.595986

**Authors:** Qiaojuan Xi, Hongkai Gao, Lan Wang-Erlandsson, Jianzhi Dong, Fabrizio Fenicia, Hubert H. G. Savenije, Markus Hrachowitz

## Abstract

Adaptation of ecosystems’ root zones to climate change critically affects drought resilience and vegetation productivity. However, a global quantitative assessment of this mechanism is missing. Therefore, we analyzed observation-based data and found that the global average root zone water storage capacity (*S*_R_) increased by 11%, from 182 to 201 mm in 1982-2020. This increase amounts to 1657 billion m^3^ over the past four decades, affecting hydrological and ecological processes worldwide. *S*_R_ increased in 9 out of 12 land cover types, while three relatively dry types experienced decreasing trends, potentially suggesting the crossing of ecosystem tipping points. Our results underscore the importance of considering root zone dynamics while analysing floods, droughts and carbon sequestration under climate change.

## Introduction

It is not possible to observe the root zone underlying the land surface at large scales directly with available observation technology. As such, it is one of the most uncertain components of the global terrestrial ecosystem ^1,2^. Climate change leads to substantial meteorological drought intensification and extension ^3,4^, which can be associated with terrestrial ecosystem degradation, mortality, or even collapse ^5,6^. However, at the same time, observations of vegetation greening indicate that, hitherto, increasing droughts have not resulted in an overall decline of global-scale vegetation^7,8^. Most studies have attributed the increase of vegetation productivity to the CO_2_ fertilization effect and land-use management ^7,8^. Nevertheless, the role of belowground root zone adaptation to drought has largely been overlooked.

The root zone water storage capacity (*S*_R_) of terrestrial ecosystems is a buffer to guarantee vegetation access to moisture during critical periods of drought ^9,10^. For a given *S*_R_, the vertical distribution of moisture may vary and its quantity does not necessarily correspond to the total volume of water in the unsaturated zone, but rather to the part of the soil moisture that is accessible to roots. For example, in the Loess Plateau where the unsaturated zone is thick, the root zone is limited to the active shallow layer of the topsoil ^11^. On the other hand, in karst and rocky substrates, the root zone may include not only the soil water storage but also the fissure water storage in the bedrock ^12^. In rainfall-runoff processes, the *S*_R_ controls the partitioning of precipitation into drainage and evaporation. As such it is a key parameter not only in hydrological models, but also in land surface models, where it controls latent and sensible heat fluxes, as well as in ecohydrological models where it regulates vegetation dynamics ^13,14^. It is well-documented that the accurate estimation of *S*_R_ significantly improves hydrological and land-surface model performance ^13-15^. Large-sample data analysis revealed that ecosystems tend to optimize their root zones to allow for the most efficient extraction of water from the substrate, thereby meeting water demand (evaporation) while minimizing their carbon expenditure for root growth and maintenance ^16,17^. *S*_R_, as an integrated reflection of root density, distribution, depth, and lateral extension can serve as an effective proxy for belowground biomass.

Ecosystems’ root zones are part of ecosystems’ survival strategy and adaptive capacity, and respond dynamically to changes in climatic and anthropogenic drivers ^18^. Such changes in *S*_R_, directly feeds back to evaporation and biomass production, thus dynamically regulating the water cycle as a whole ^9,19^. Globally, *S*_R_ varies across landscapes and climatic zones, with larger *S*_R_ in areas with high seasonality, high aridity, longer dry periods, and larger rainfall variability (for example, tropical savanna and Mediterranean climate), and smaller *S*_R_ under relatively stable conditions (for example, tropical rainforests and temperate climates). Human activities, such as deforestation and irrigation, can also alter *S*_R_. In experimental catchments in the US and Germany, *S*_R_ was observed to sharply decline immediately after deforestation, and then gradually recover over a period of 5∼13 years ^20^. Irrigation on agricultural land providing additional water during dry periods, artificially reducing the exposure of vegetation to droughts, leading to a smaller *S*_R_ than under natural conditions ^10^.

Although *S*_R_ is a key variable in runoff generation, drought resilience and land-atmosphere interactions, a global quantitative assessment of the temporal evolution and spatial variation of *S*_R_ in response to climate change and anthropogenic activities is missing. Classical approaches to estimate *S*_R_ are based on combining information of soil texture and rooting depth from field observations using look-up tables by land-use categories, which are problematic to upscale. More importantly, being mostly snapshots in time, these approaches ignore the temporal dynamics of *S*_R_ ^21^. Inverse model approaches based on satellite data of precipitation, evaporation and other meteorological factors have previously been shown to be valuable to estimate rooting depths at large scales. However, such approaches require extensive parametrizations and assumptions on soil-plant-water relationships ^16,22^ Some terrestrial biosphere models now use aboveground biomass changes as a surrogate of belowground responses ^23^. However, the underlying assumption of a stationary roots-shoots ratio remains to be tested.

In contrast to these methods, the mass curve technique (MCT), initially developed for determining reservoir capacity in civil engineering ^24^, allows for model- and scale-independent estimates of the *S*_R_. The MCT approach makes use of the water balance and merely requires cumulative vertical inflow (rainfall, snowmelt, and irrigation) and outflow (dry spell evaporation) ^9^, which are all observable variables from e.g., in-situ measurements, reanalysis data, and satellite remote sensing ^10^. Such time series of inflows and outflows are used to infer water deficit during dry spells, where the largest deficit over a sufficiently long period of time can be assumed to represent the actual *S*_R_. The MCT approach has previously been used to estimate present-day *S*_R_ at both the catchment and global scale, and shown to yield similar results as the inverse modelling approach and outcompete the look-up table approach ^10,19^. Nevertheless, the MCT approach has yet to be used for quantifying global *S*_R_ dynamics in a changing environment.

In this study, we aim to quantify the temporal trend and spatial variation of *S*_R_, identify the drivers of observed change, and explore the contributions of *S*_R_ change to global greening. We applied the MCT approach to state-of-the-art high-quality reanalysis data to quantify *S*_R_ from 1982 to 2020 at the grid cell, regional, and global scales. Furthermore, we investigated the effect of drought duration and average daily water deficit on *S*_R_ trends. To explore the adaptation of global ecosystems to drought, we analyzed the relationship between belowground *S*_R_ and aboveground Leaf Area Index (LAI) changes at grid cell and regional scales. Finally, we used *S*_R_ to infer the change of belowground biomass over the past four decades.

## Results

### Increasing trend of global *S*_R_ with increasing drought duration

The global average annual *S*_R_ increased from 182 mm to 201 mm in the period of 1982-2020, with a trend of 0.57 mm yr^-1^ (*p* < 0.001; Fig.1). Multiplied by the terrestrial vegetation area (85 million km^2^), the total increased volume of *S*_R_ over this period is 1657 billion m^3^, equivalent to the total storage capacity of 42 Three Gorges reservoirs, the largest hydraulic engineering project in the world. Most of the increase in *S*_R_ took place 1991-2007. In 1982-1990, *S*_R_ in fact decreased, and in 2008-2020, *S*_R_ reached a plateau (Fig.1).

**Fig. 1.**
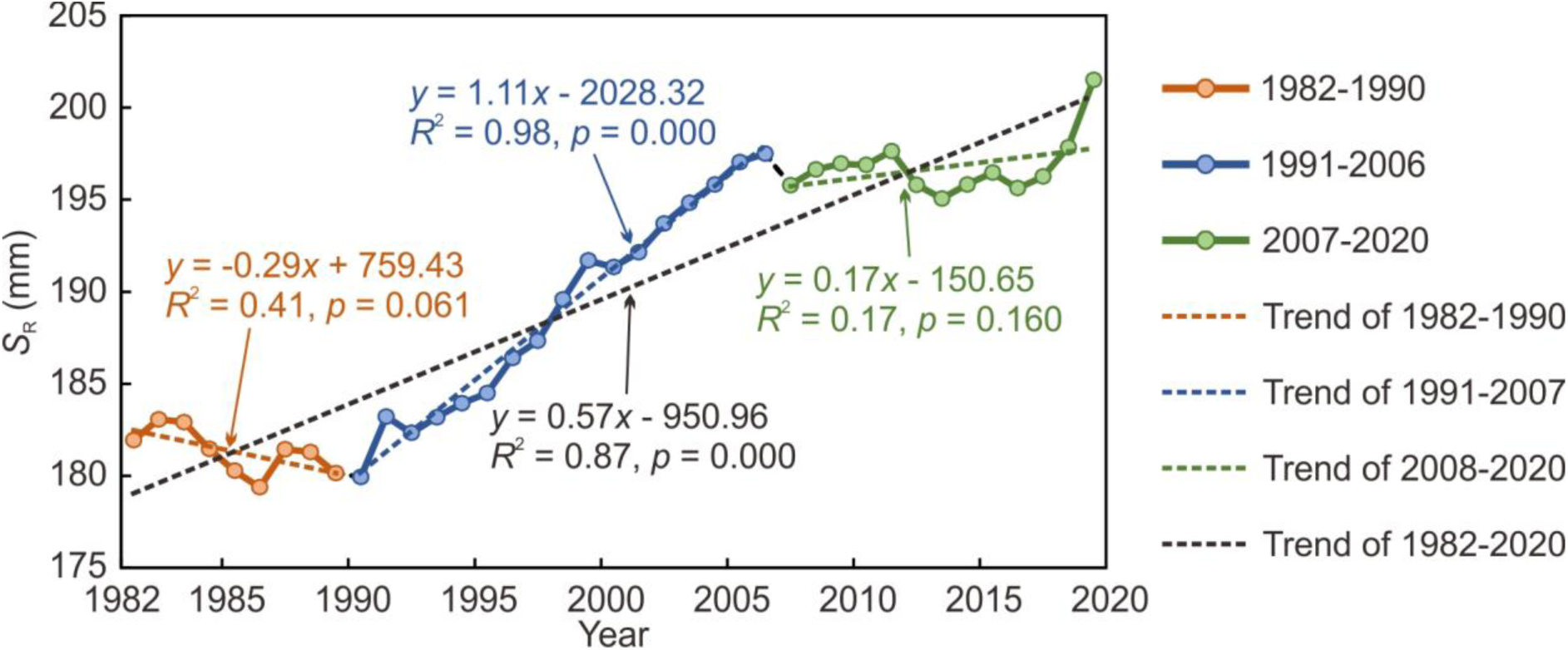
Trends of global average *S*_R_ during the period 1982 to 2020. The dots represent the values of *S*_R_ for each year.

We found a concurrent increasing trend in both *S*_R_ and the aridity index (AI) (Fig. 1; Extended Data Fig. 1). To further explore how drought affects *S*_R_, we decomposed *S*_R_ change into the changes due to drought duration (*L* in days) and average daily water deficit during drought (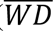 in mm d^-1^) (see Methods; Supplementary Fig. S1b). Overall, global *S*_R_ increase was found to be significantly influenced by increasing drought duration (Extended Data Fig. 2). However, the contributions of drought duration and average daily water deficit vary across different time periods, with average daily water deficit driving *S*_R_ changes in the first period (1982-1990) and drought duration playing a more prominent role in the recent period (1991-2020; Extended Data Fig. 2).

### Spatial pattern of *S*_R_ variation and its drivers

The changes in *S*_R_ are spatially heterogeneous. At the grid cell scale, we observed increasing *S*_R_ trends in 66% of the global vegetated land (54%, *p* < 0.1; 12%, *p* ≥ 0.1), and decreasing trends in 34% (24%, *p* < 0.1; 10%, *p* ≥ 0.1; see Fig. 2a; Extended Data Fig. 3). Increasing trends are common in central US, central Africa, northern Eurasia, and central South America, whereas decreasing trends are mostly found in dry regions of the western US, southern Asia, northern China, southern Africa, and northeastern Australia.

**Fig. 2.**
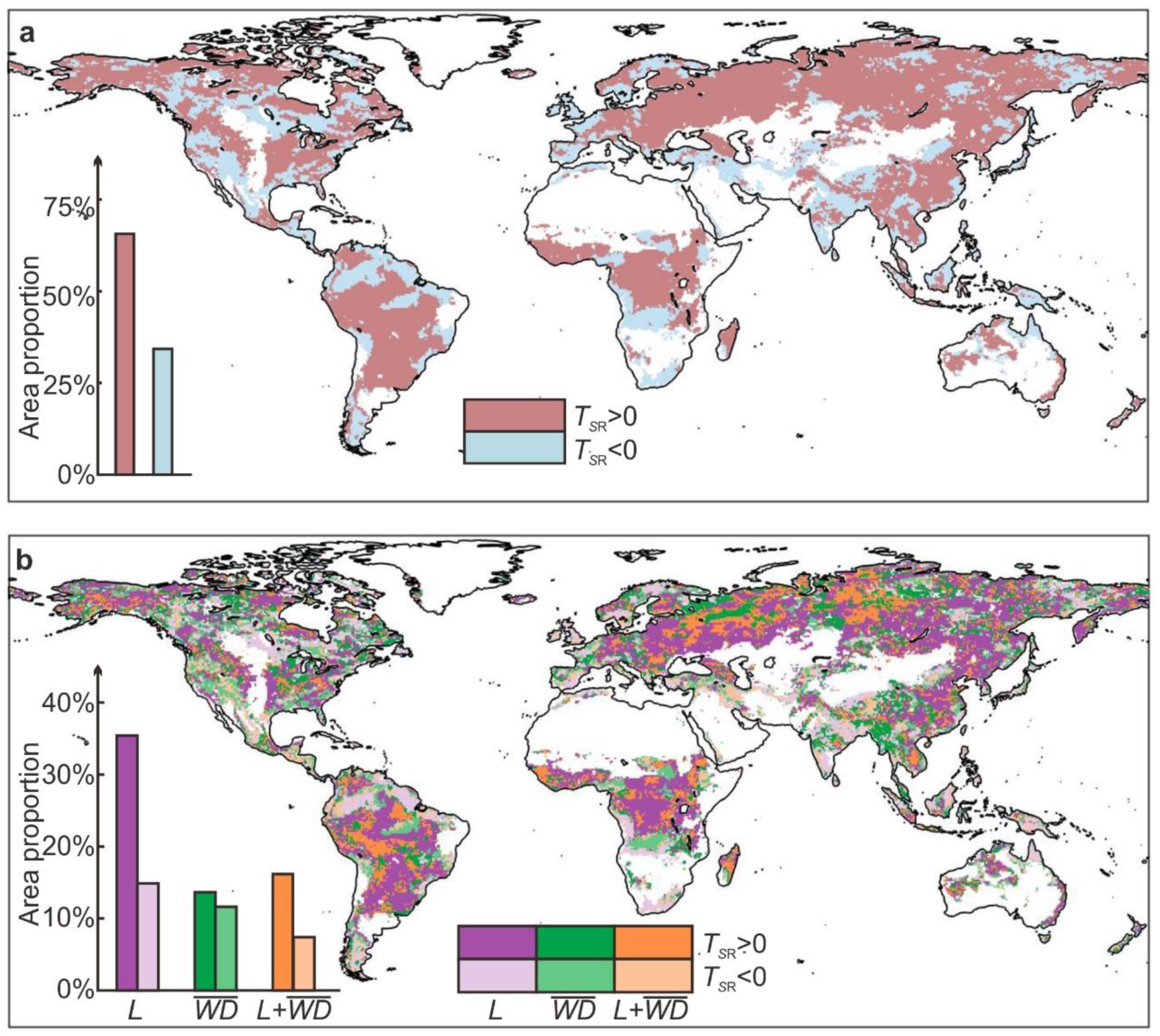
Spatial pattern of *S*_R_ changes and its drivers. Panel (**a**): spatial distribution of the direction of *S*_R_ trends 1982-2020. *T_S_*_R_ denotes the trend of *S*_R_. The histogram shows the percentage of areas with different *S*_R_ change types across global vegetated land (total 85 million km^2^). Panel (**b**): dominant drivers affecting *S*_R_ change at grid cell scale. 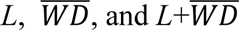 denote drought duration, average daily water deficit, and both factors, respectively. The histogram shows the percentage of areas with different dominant factors for *S*_R_ changes across global vegetated land.

We also examined the drivers of *S*_R_ change for each grid cell. Of the land areas with increasing *S*_R_, 54% was associated with an increase of drought duration, 21% with a rise in average daily water deficit, and 25% with an increase of both (Fig. 2b). Of the land areas with decreasing *S*_R_, we found diminishing proportions of influence from drought duration, average daily water deficit, and the combined effect of both factors, accounting for 44%, 34%, and 22%, respectively (Fig. 2b). In summary, changes in drought duration play a dominant role in *S*_R_ dynamics.

Finally, we analyzed *S*_R_ trends in 12 land cover types and found increasing *S*_R_ trends in 9 types and decreasing trends in 3 relatively dry types (Fig. 3). Increasing trends were found in forests, savannas, grasslands, and croplands, among which deciduous broadleaf forests has the largest increasing trend (1.10 mm yr^-1^), followed by mixed forests (0.93 mm yr^-1^). Decreasing trends were found in shrublands and barren land, with closed shrublands experiencing the largest decrease (0.94 mm yr^-1^). Further, we explored the drivers of *S*_R_ change in different land cover types. *S*_R_ increase in most types were mainly associated with the increase in drought duration, except for mixed forests which was driven by the increase of both drought duration and average daily water deficit (Extended Data Fig. 4). *S*_R_ decrease in shrublands was mainly associated by a decline in average daily water deficit, and in barren land with the decrease in drought duration (Extended Data Fig. 4).

**Fig. 3.**
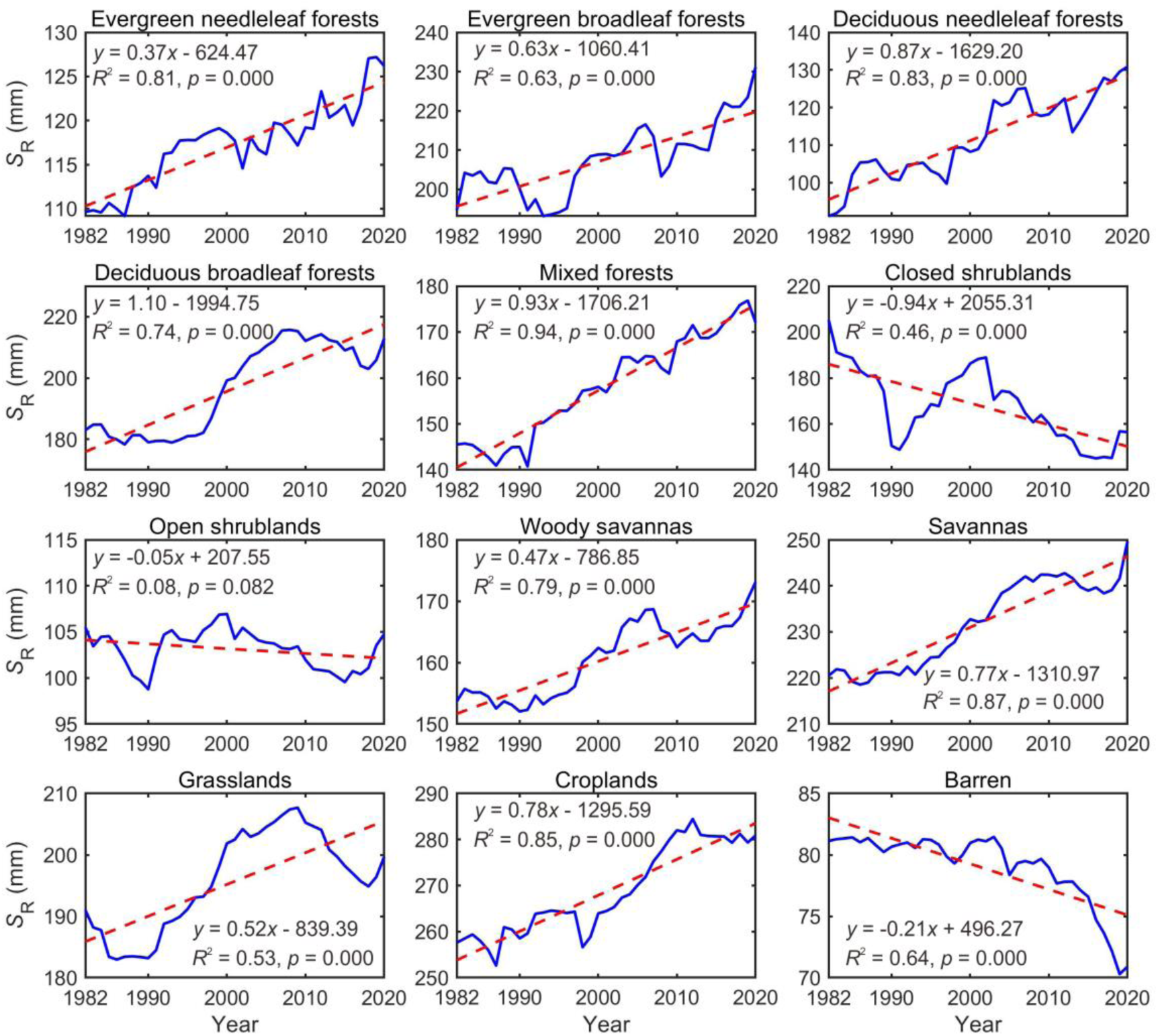
Change of *S*_R_ in different land cover types in 1982-2020.

### Comparison between belowground *S*_R_ and aboveground greenness

We compared the changes of belowground *S*_R_ with aboveground greenness (as represented by LAI) at the grid cell and regional scale to explore ecosystems’ adaptation strategies to climate change. During 1982-2020, both *S*_R_ and LAI increased in a considerable fraction of global vegetated land (56%), *S*_R_ increased while LAI decreased in 10%, *S*_R_ and LAI decreased simultaneously in 5%, and *S*_R_ decreased while LAI increased in 29% of vegetated land (Fig. 4). Concurrent increasing trends of both *S*_R_ and LAI are mainly found in central US, the northern Amazon rainforest, eastern Europe, Siberia, eastern Asia, and the Congo rainforest, which are mostly in drying regions (Extended Data Fig. 5). This can be interpreted as vegetation in these areas enhancing *S*_R_ to adapt to increasing droughts while sustaining aboveground greening. Concurrent *S*_R_ increase and LAI decrease were mainly concentrated in boreal and tundra regions of northern North America and northern Asia, implying that the resilience of these ecosystems was threatened by climate drying, manifested as an inability to sustain aboveground productivity increase. Areas with both *S*_R_ and LAI decreasing are distributed across the world, but are more common in the pan-Arctic and arid regions. These ecosystems might already be transitioning or collapsing due to drought-driven stress. Decreasing *S*_R_ trends that coincide with increasing LAI trends mainly appear in regions of extreme humidity (such as northern South America) and intensive human activities (such as western Europe, India), and might be explained by climate wetting (Extended Data Fig. 5) and/or human induced land cover change (such as agricultural expansion and/or intensification) ^2,8,25^.

**Fig. 4.**
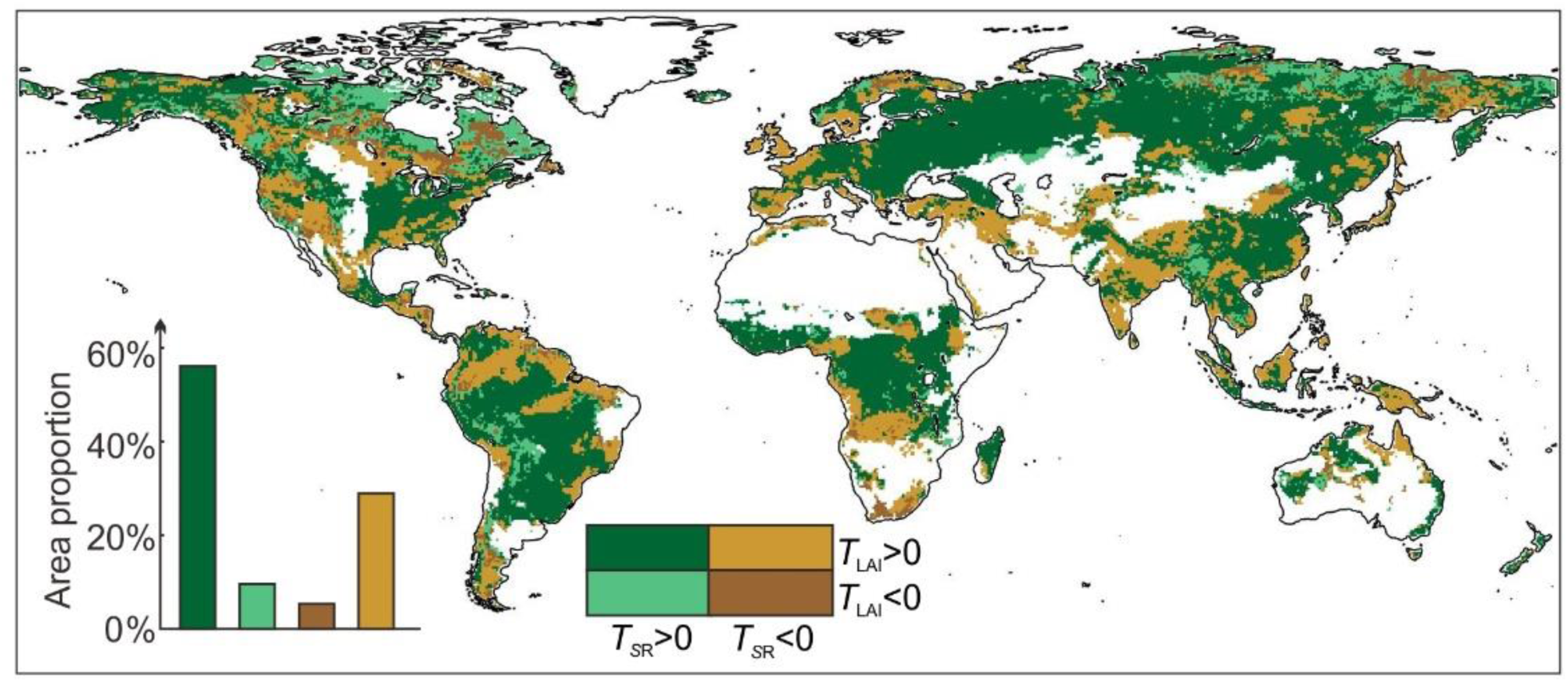
The comparison between *S*_R_ and LAI from 1982 to 2020. The spatial pattern illustrates the areas with different colors for the four combinations of positive/negative *T_S_*_R_ (Trend of *S*_R_) and *T*_LAI_ (Trend of LAI) in 1982-2020. The white regions represent areas with no vegetation cover and high uncertainty data. The accompanying histogram demonstrates the proportion of different combinations in the global vegetated area.

As for different land cover types, *S*_R_ and LAI simultaneously increased in 9 out of 12 types (Fig. 3; Extended Data Fig. 6), suggesting ecosystem in most land cover types enhanced root zone to adapt to intensified drought (Extended Data Fig. 7). In 3 relatively dry land cover types, *S*_R_ decreased with increasing drought, while LAI showed a weakly increasing trend (Fig. 3; Extended Data Fig. 6; Extended Data Fig. 7). Positive effects of CO_2_ fertilization and rising temperature on greening of these cover types may offset negative effects of *S*_R_ decrease. LAI increase in closed shrublands and barren land may be mainly attributed to CO_2_ fertilization ^7^, whereas the greening in open shrublands may be mainly attributed to rising temperatures, enhanced photosynthesis and longer growing seasons ^7^.

### Increase of belowground biomass

To estimate belowground biomass change, we assumed a positive linear proportional relationship between *S*_R_ and belowground biomass. This assumption ignores non-linear interactions between root zone water, carbon, nutrient, and microbial processes, but can be valid in water-limited shrublands and grasslands where increases in belowground biomass are dominated by water availability ^26,27^. In humid forests, the assumption can lead to under or overestimation in belowground biomass, since the changes may be affected by nutrition, and because the fine roots mainly used for water uptake have lower biomass than coarse roots ^28^. The baseline for belowground biomass in global and land cover types scales was taken from Ma et al. ^29^, and belowground biomass changes were estimated based on *S*_R_ change (see Methods).

The analyses suggest a global increase in belowground biomass of 12.2 (9.7∼14.6) GtC from 1982 to 2020. Among the different land cover types, the largest increase in total and unit area was found in evergreen broadleaf forests (3.9 GtC in total, 3.5 tC ha^-1^ in per unit area; Extended Data Fig. 8); while the largest total reduction was found in open shrublands (0.1 GtC) and largest unit area reduction in closed shrublands (2.1 tC ha^-1^; Extended Data Fig. 8).

## Discussion

This study shows a global increasing trend in *S*_R_ over the past four decades caused by increasing drought (primarily driven by drought duration, but also by the average daily water deficit during drought) reflected in the aridity index (AI). Intensifying drought increases ecosystems water demand, as a result of which dominate ecosystems allocated more carbon and nutrients to enhance their root zone systems^30^, leading to an increase in *S*_R_ to sustain its water supply. We found that global *S*_R_ slightly decreased from 1982 to 1990, then sharply increased until around 2007, after which it levelled off. Notably, *S*_R_ decreased abruptly in shrublands and barren land (Fig. 3), while LAI showed a slight increasing trend (Extended Data Fig. 6). But the increase of aboveground biomass (LAI) does not mean that ecosystems experience an increasing resilience, which may lead to an expanding but more vulnerable ecosystem risk ^31^. This potentially implies that ecosystems in these land cover types may have crossed ecosystems’ thresholds or tipping points, which aligns with the breakpoints detected in global arid ecosystems using rain-use efficiency ^32^ and Normalized Difference Vegetation Index (NDVI) in remote sensing observations ^33^. Additionally, the decreasing *S*_R_ in arid regions shortens the response time to drought ^34^, consistent with reports of degradation and regime shifts in arid grasslands, deserts, and drylands globally ^32^.

We found that most terrestrial ecosystems have tended to increase their root zone to adapt to intensifying droughts, and simultaneously sustaining Earth’s greening in the past four decades. The ways in which ecosystems adapt or respond to changing climatic conditions are diverse. For instance, if there is no shortage of water by aboveground greening due to the CO_2_ fertilization, but alternatively by increasing *S*_R_, or by ecosystem transition (e.g. forest to savannah, or savannah to dry land) ^35^, or by shifting in microbial communities ^36^, to cope with increasing droughts. Our finding implies that increasing *S*_R_ is likely one of the most effective and swift responses of ecosystems to increasing drought. In regions with increasing *S*_R_, ecosystems are still adapting to a changing climate. In regions with decreasing *S*_R_, ecosystems may have reached the limit of their adaptive capacity. Neglecting *S*_R_ change in impact studies can, thus, lead to both under- and overestimations of climate risks to ecosystem health ^35,37^.

As an important part of the underground carbon pool, the root zone is critical for the global carbon budget. Our study shows that an increase in belowground biomass carbon may be significant in evergreen broadleaf forests, further highlighting the importance of tropical ecosystems in the global carbon sink ^38^. The plant biomass allocation to leaves, stems, and roots has commonly been assumed to be a constant value ^29^, but without rigorous testing. The roots-shoots ratio is a vital expression to capture biomass allocation in plants, since shoots and roots are generally the resource-acquiring organs in comparison with stems ^39^. Here, we found that while changes in roots and shoots – here loosely interpreted as trends in *S*_R_ and LAI, respectively, have the same trend in 61% of vegetated land areas, changes in the opposite directions are found in 39% of the land. Hence, our results do not support the assumption that plant biomass allocation remains constant under climate change. Globally, we found a higher increase in belowground root zone (11%; Fig.1) concurring with aboveground greening (6%; Extended Data Fig. 9).

The ERA5 evaporation data employed for deriving *S*_R_ is expected to reproduce realistic evaporation dynamics, since the model-simulated evaporation from the land surface model HTESSEL used in the ERA5 ^15,40^ is adjusted based on assimilation using remotely sensed soil moisture and near-surface atmosphere conditions (e.g., air temperature and specific humidity) ^41^. It has been shown that the assimilation procedure is able to correct evaporation for irrigation effects, even when irrigation is not applied in the land surface model ^42^. In addition, our results are also quantitively confirmed by the FLUXCOM global evaporation dataset, which is based on an entirely data-driven algorithm (Fig. 5a).

**Fig. 5.**
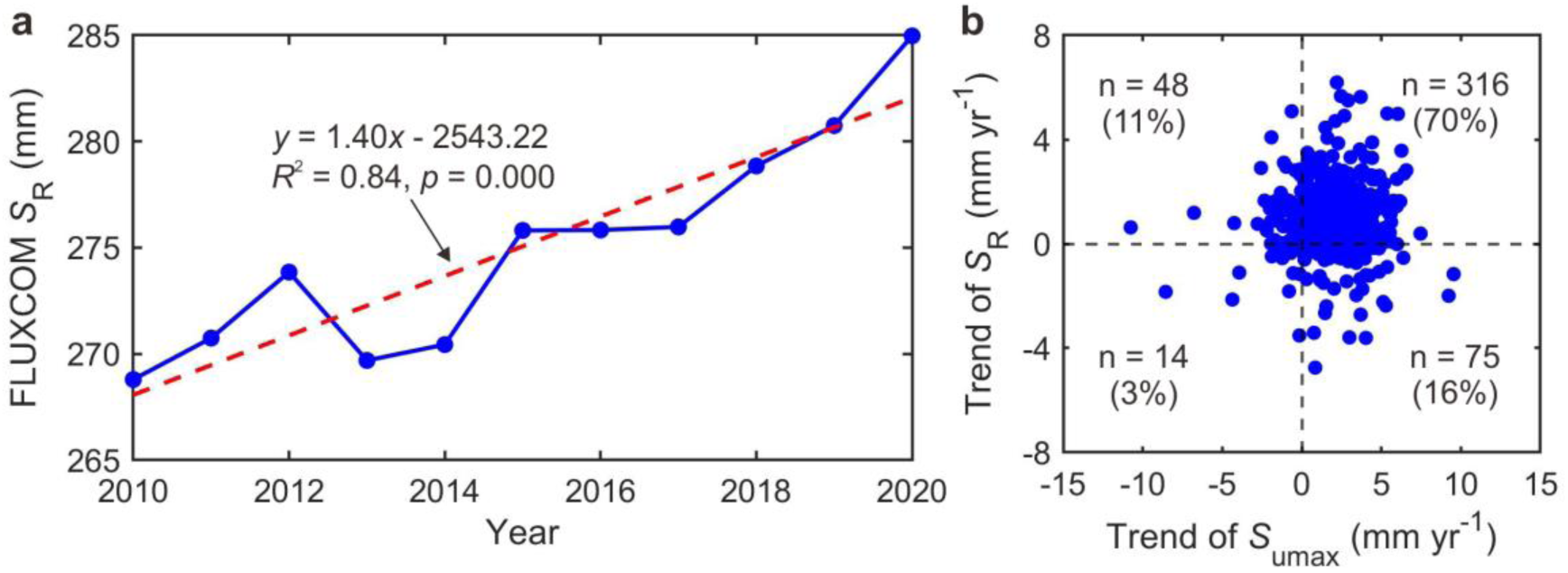
More independent evidences supporting the increasing trend of root zone water storage capacity. (**a**) The temporal trend of *S*_R_ estimated by FLUXCOM in 2010-2020. (**b**) The scatter comparison of the *S*_R_ and *S*_umax_ trend of 453 CAMELS catchments in the USA. One dot represents one catchment.

More importantly, basin-scale water balances also support our findings that *S*_R_ increases under climate change. Based on long-term streamflow records from 453 CAMELS catchments in the US ^43^, we calibrated the root zone water storage capacity parameter *S*_umax_ in a standard process-based hydrological model (FLEX) ^44,45^, by the Dynamic Identification Analysis (DYNIA) method ^46,47^. We found that 86% (391 out of 453) catchments demonstrated an increasing trend of *S*_umax_, which provides a strong and independent evidence to support our main findings (Fig. 5b).

Finally, a meta-analysis from field experiments across 110 published studies indicated that vegetation expands its root systems globally in response to the elevated concentration of CO_2_ ^48^, which agrees with our result of global *S*_R_ increase. The increasing trend of *S*_R_ is also consistent with globally increasing soil moisture deficits ^49^, as vegetation water use reduces soil moisture and increases *S*_R_. Moreover, the global annual fluctuation of *S*_R_ and total water storage (TWS) variability from Gravity Recovery and Climate Experiment (GRACE) data showed a positive correlation (Extended Data Fig. 10), which provides another evidence showing the synchronicity of the increasing of terrestrial total water deficit and increase of *S*_R_, as the maximum deficit of root zone water storage. In summary, our finding of an increasing trend in root zone water storage capacity is supported by multi-source data (ERA5, CAMELS, FLUXCOM, and GRACE) and independent methods (MCT and DYNIA).

Despite use of state-of-the-art methods and datasets, and alignment with independent sources, the estimated *S*_R_ is still subject to uncertainty associated with forcing data quality. The method of estimating belowground carbon still needs further refinement and should be tested by observations with controlled experiments or field studies. Combined with an asymptotic equation ^50^, *S*_R_ may be used to estimate fine root carbon. In this study, we implicitly considered the effect of CO_2_, as both vegetation and hydrological processes are affected by CO_2_, temperature, and precipitation. However, isolating the specific effect of CO_2_ on *S*_R_ is difficult. Moreover, the MCT is a diagnostic model rather than a prognostic model. Future studies need to consider the full effects of climate change on vegetation physiology to project *S*_R_ variation in future scenarios.

To the best of our knowledge, this is the first study to quantify the spatio-temporal variation of *S*_R_ at global scale. We believe this study has improved our understanding of the mechanism of terrestrial ecosystems’ resilience to drought and the role of belowground adaptation in maintaining the terrestrial biosphere within a safe operating space for humanity.

## Methods

### Mass curve technique (MCT)

The mass curve technique (MCT), a deficit-based approach, was adopted to estimate *S*_R_ ^9,10,51,52^. The method is based on the water balance. Inflow (*F*_*in*_) to the root zone is the sum of precipitation and snowmelt and irrigation, and outflow (*F*_*out*_) is evaporation. This approach considers the daily evaporation, precipitation, snowmelt, and temperature data from the ERA5 reanalysis product at a spatial resolution of 0.5°for the period 1971-2020 (https://cds.climate.copernicus.eu/cdsapp#!/software/app-c3s-daily-era5-statistics?tab=app). To determine whether precipitation was rainfall or snowfall, we used the daily temperature. If the temperature was below 0°C, precipitation was considered snowfall, and its value of rainfall for that day was set to zero. For irrigation data from 1971 to 2010, we used an average of four model outputs (WaterGAP, H80, LPJmL, and PCR-GLOBWB) adjusted by correction factors between model estimates and reported estimates ^53^, with a monthly temporal resolution and 0.5°spatial resolution. For the period from 2011 to 2020, we assumed that irrigation remained unchanged compared to 2010. We downscaled the monthly irrigation data to daily resolution based on ERA5 daily evaporation.

The estimation of *S*_R_ involves calculating the water deficit in the root zone when the outflow (evaporation) exceeds the inflow (the sum of precipitation, snowmelt, and irrigation). This deficit indicates that plants need to use the water stored in *S*_R_ to sustain their water use, such as transpiration. The calculation of the outflow and inflow water deficit for each day is performed as follows (Equation 1):

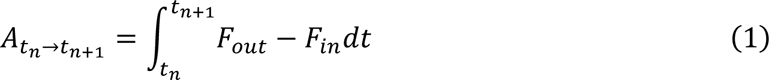

*A*_*tn*→*tn*+1_ is the water deficit in *t*_n+1_ day. The aggregate of water deficit in each day is the accumulative water demand (Equation 2).

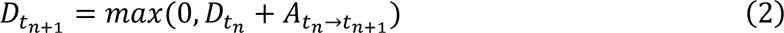

*D*_*tn*+1_ is the accumulative water deficit in day *t*_n+1_. The dry period varies for different climates. For example, the dry period of monsoon climate is mainly in winter, while the dry period of Mediterranean climate is mainly in summer. Considering these seasonality variations among different climates, *D* does not get back up to zero and the deficit is carried over to the next year. In general, the inflow of the root zone exceeds the outflow for annual totals. Therefore, we removed grid cells where the average annual inflow was less than outflow from 1971 to 2020 so as to exclude the influence of abnormal data. But in very rare cases, such as in arid regions, *D* may also accumulate over more than one year and data were reset if the accumulation extended over two consecutive years. Note that *D* never becomes negative, as it can be considered a running estimate of the root zone storage reservoir.

Since the *S*_R_ is the result of ecosystems’ relative long-term response to climate ^9^, we select the maximum of a number of years’ *D* as the *S*_R_ of the study year:

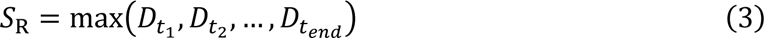

Because earlier work has shown that globally ecosystems tune to a drought of once in about 10 years ^9^, we used the maximum *D* of the past ten years as the *S*_R_. For example, the maximum *D* from 1973 to 1982 is referred to as the *S*_R_ of 1982. Here, we used data from 1971 to 1972 as a spin-up period to prevent abnormal initial conditions. The conceptual illustration of the algorithm and some examples at the grid cell scale for calculating *S*_R_ are shown in Supplementary Figs. S1a and S2.

### Trend analysis

In this study, we conducted an analysis of the temporal and spatial patterns of *S*_R_ (root zone water storage capacity) in global vegetated lands at different scales. For the analysis of the global *S*_R_ change trend, we used regression analysis to estimate the trend of average *S*_R_. The average *S*_R_ was calculated by area-weighting, since the area of 0.5°grid is not equal across latitudes. In order to analyze *S*_R_ change at the regional scale, we referred to MODIS Land Cover Type product (MCD12Q1) (https://lpdaac.usgs.gov/products/mcd12q1v006/) at 500 m spatial resolution for 2020 to define land cover types. The land cover type product was resampled to 0.5°using the majority algorithm which determines the new value of the cell based on the most popular values in the filter window. We chose 12 land cover types (Supplementary Fig. S3), including evergreen needleleaf forests, evergreen broadleaf forests, deciduous needleleaf forests, deciduous broadleaf forests, mixed forests, closed shrublands, open shrublands, woody savannas, savannas, grasslands, croplands, and barren, to calculate the trend of area-weighting average *S*_R_ by regression analysis. The *S*_R_ trend for each grid cell was also calculated by regression analysis.

To further explore how plants adapt to climate change, we analyzed the trend of leaf area index (LAI) and arid index (AI) at the grid cell and global scale. The LAI data used in our study was obtained from the GIMMS LAI4g dataset which utilized spatiotemporally consistent BPNN models and the latest PKU GIMMS NDVI product and high-quality global Landsat LAI samples to remove the effects of satellite orbital drift and sensor degradation ^54^. The GIMMS LAI4g dataset provided half-month temporal resolution for the period 1982-2020, with a spatial resolution of 1/12°. Abnormal values exceeding three times the maximum value of the three preceding and following temporal data points or falling below one third of the minimum value were removed from the dataset. To fill missing values in the LAI temporal series, linear interpolation was used, and Savitzky-Golay filtering was applied to smooth the data using a window width of 7 and a smoothing polynomial of 3. The LAI data were resampled to a coarser resolution of 0.5°, and the annual average LAI was calculated for further analysis. We conducted regression analysis to calculate trends of LAI at the grid cell and global scale from 1982 to 2020.

The AI for the period 1982-2020 in this study was calculated through annual potential evaporation and precipitation using Equation 4:

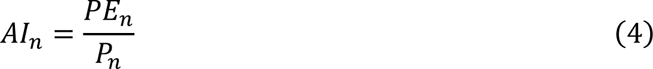

where *AI*_*n*_ is the aridity index in year *n*, *PE*_*n*_ is the annual potential evaporation in year *n*, *P*_*n*_ is the annual precipitation in year *n*. The potential evaporation was from Global Land Evaporation Amsterdam Model (https://www.gleam.eu/) at daily temporal resolution for 1982-2020 with 0.25°spatial resolution. We resampled potential evaporation data to 0.5°, and calculated annual potential evaporation at the global and grid cell scale. Annual precipitation was calculated from ERA5 daily precipitation. The AI trend for the global and grid cell scale was calculated by using regression analysis.

### Attribution analysis

The MCT method also provides a powerful diagnostic tool to gain valuable insights into the drivers of *S*_R_ changes and their implications for ecosystem response to climate change and water availability. Based on the MCT method, *S*_R_ represents the maximum accumulation of water deficit in the root zone during drought periods. To explore the factors driving *S*_R_ change, we calculated the days corresponding to the *S*_R_ accumulated period and determined the average daily water deficit by calculating the average difference between outflow (evaporation) and inflow (precipitation, snowmelt, and irrigation) during that period. Using this approach, we decomposed *S*_R_ into two main components: drought duration and average daily water deficit during drought, where *S*_R_ is equal to the product of drought duration and average daily water deficit (Supplementary Fig. S1; Supplementary Fig. S2).

In this study, we investigated the impact of two drivers on *S*_R_ from 1982 to 2020 at the grid cell, regional, and global scales, and these two drivers correspond one-to-one with *S*_R_. Based on this decomposition, if *S*_R_ increases, it must be caused by an increase in drought duration, average daily water deficit, or both; and vice versa. We categorized the factors driving *S*_R_ changes into three main categories (Supplementary Fig. S1b): (1) If the direction of the *S*_R_ trend is the same as drought duration but opposite to average daily water deficit, the *S*_R_ change is considered to be dominated by changes in drought duration. (2) If the direction of the *S*_R_ trend is inverse to drought duration but the same as average daily water deficit, the *S*_R_ change is considered to be dominated by changes in average daily water deficit. (3) If the direction of the *S*_R_ trend coincides with both drought duration and average daily water deficit, changes in both factors are considered to contribute to the change in *S*_R_.

Using regression analysis, we calculated drought duration and average daily water deficit trends at the grid cell, regional, and global scales, and determined the respective contributions of the two to the changes in *S*_R_. And the trends between drought duration and average daily water deficit are independent. This allowed us to understand the relative importance of these two factors in driving the observed changes in *S*_R_ over time.

### Comparison between *S*_R_ and LAI dynamics

To compare the changes in *S*_R_ and LAI from 1982 to 2020, we overlaid the regression trend of *S*_R_ (*T_S_*_R_) and LAI (*T*_LAI_) to analyze the differences between their trends. The comparison of *S*_R_ and LAI trends can be divided into four categories: *T_S_*_R_>0 & *T*_LAI_>0, *T_S_*_R_>0 & *T*_LAI_<0, *T_S_*_R_<0 & *T*_LAI_>0, and *T_S_*_R_<0 & *T*_LAI_<0.

### Estimation of changes in belowground biomass

The belowground biomass data used in this study were obtained from Ma et al. ^29^ and derived by combining aboveground biomass data with the root-shoot ratio. The original data had a spatial resolution of 0.0083°, which we resampled to a coarser resolution of 0.5°. In this study, we assumed that the change in belowground biomass is proportional to the change in *S*_R_. We took the underground biomass data from Ma et al. ^29^ as the belowground biomass data in 2020 and calculated belowground biomass in different land cover types by area-weighting. Then, the proportion of *S*_R_ change from 1982 to 2020 relative to 2020 was computed according to the *S*_R_ regression trend in different land cover types using Equation 5, 6 and 7:

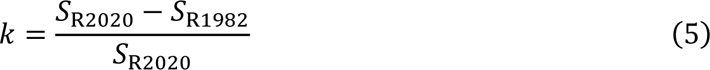

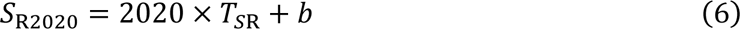

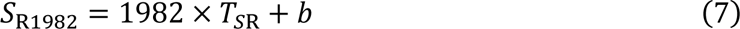

where *k* is the proportion of *S*_R_ change, *T_S_*_R_ and *b* represent the regression trend and intercept of *S*_R_ from 1982 to 2020 respectively, *S*_R2020_ and *S*_R1982_ are the *S*_R_ values for year 2020 and 1982 calculated by the regression equation. Last, belowground biomass in 2020 was multiplied by the change proportion of *S*_R_ to estimate the change from 1982 to 2020 (Equation 8):

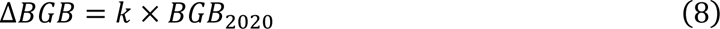

where Δ*BGB* is the change of belowground biomass in 1982-2020, *BGB*_2020_ is the belowground biomass in 2020. It should be noted that although this is a reasonable assumption, we did not consider exponential decay of root biomass with depth and possibly overestimated the change of belowground biomass.

### Uncertainty analysis

To further test the robustness of our findings on *S*_R_, we estimated *S*_R_ using FLUXCOM from 2001 to 2020 (https://www.bgc-jena.mpg.de/geodb/projects/Home.php) as input evaporation at the grid cell scale. The FLUXCOM was produced based on machine learning methods that integrate FLUXNET site level observations, satellite remote sensing and meteorological data. The FLUXCOM data from 2001 to 2015 has a monthly temporal resolution and 0.5°spatial resolution, while the data from 2016 to 2020 has 8-day temporal resolution and 0.083°spatial resolution. The FLUXCOM data was resampled to 0.5°, and downscaled to daily temporal resolution referring to ERA5 daily temperature. We used the MCT method to estimate *S*_R_ and analyzed the trend of the global average *S*_R_ using the maximum values over a ten-year period.

Another approach for estimating root zone water storage capacity is by calibration of a hydrological model to streamflow observations. The model used in this study was the FLEX hydrological model which simulates hydrological processes at catchment scale ^44,55^. There are 10 free parameters in the FLEX model that require calibration. Among them, *S*_umax_ represents the root zone water storage capacity. The input to the model was the CAMELS (Catchment Attributes and MEteorology for Large-sample Studies) dataset ^43^. The CAMELS dataset provides daily meteorological forcing (including precipitation and temperature) and discharge for catchments across the United States from 1980 to 2014. The Dynamic Identification Analysis method (DYNIA) was used to assess the temporal variation of the model parameters ^46^. In this study, we generated 40,000 sets of parameter combinations within the feasible range for 10 parameters, basing on a Monte Carlo framework and employing a Latin hypercube sampling technique. Each set of parameters is associated to a streamflow simulation, for which a performance metric is calculated by the Kling-Gupta efficiency (KGE) using a five-year moving window ^47^. We selected the optimal parameters with highest KGE for each period and each catchment and compared the trend between *S*_umax_ and *S*_R_ in 453 catchments.

Total water storage (TWS) variability, the difference between the maximum and minimum value in a year, was also used to verify the robustness of *S*_R_. The long-term TWS was reconstructed from GRACE by combining machine learning with time series decomposition and statistical decomposition techniques for the period 1982-2019, with a monthly temporal resolution and 0.5°spatial resolution ^56^. We calculated global average TWS variability by area-weighting for 1982-2019, and then analyzed the Spearman rank correlation between *S*_R_ and TWS variability. The glacier region (https://nsidc.org/data/nsidc-0770/versions/6) and the area irrigated with groundwater over 80% (https://www.fao.org/aquastat/zh/geospatial-information/global-maps-irrigated-areas/latest-version) were excluded from this analysis, since *S*_R_ mainly considers the change of subsurface water deficit.

## Acknowledgments

H.G. acknowledges the support from the National Natural Science Foundation of China (grant nos. 42122002 and 42071081). L. W. acknowledges financial support from the European Research Council (ERC-2016-ADG-743080, Horizon Europe 101081661), Formas (2022-02089; 2019-01220), and the IKEA Foundation.

## Author contributions

Conceptualization: H.G.; Methodology: Q.X., H.G., M.H. and L.W.; Critical insights: L.W., H.H.G.S., J.D., M.H. and F.F.; Writing – original draft: Q.X. and H.G.; Writing – review and editing: H.G., Q.X., L.W., J.D., F.F., H.H.G.S. and M.H.

## Data availability

The meteorological data used in this study were publicly available from the ERA5 reanalysis product (https://cds.climate.copernicus.eu/cdsapp#!/software/app-c3s-daily-era5-statistics?tab=app). The irrigation data were sourced from the global gridded monthly sectoral water use dataset (https://zenodo.org/record/1209296#.Y2TrHWlByUk). For potential evaporation, we used data from the Global Land Evaporation Amsterdam Model (https://www.gleam.eu/). The LAI data were obtained from GIMMS LAI4g (https://zenodo.org/record/7649108). To classify land cover types, we utilized data from MODIS Land Cover Type Product (MCD12Q1) (https://lpdaac.usgs.gov/products/mcd12q1v006/). The belowground biomass data were calculated by Ma et al. (https://figshare.com/authors/Haozhi_Ma/11306118). FLUXCOM energy fluxes data were accessible from the Data Portal of the Max Planck Institute for Biogeochemistry (https://www.bgc-jena.mpg.de/geodb/projects/Home.php). The long-term TWS data were reconstructed by Li et al. (https://datadryad.org/stash/dataset/doi:10.5061/dryad.z612jm6bt). The glacier outlines (excluding the ice sheets in Greenland and Antarctica) were available from Randolph Glacier Inventory 6.0 (https://nsidc.org/data/nsidc-0770/versions/6), and the irrigation areas were obtained from Global Map of Irrigation Areas version 5 (https://www.fao.org/aquastat/zh/geospatial-information/global-maps-irrigated-areas/latest-version). The hydrological model inputs were publicly available from CAMELS (https://doi.org/10.5065/D6MW2F4D).

