## Supplement materials for "Terrestrial ecosystems enhance root zones in response to intensified drought"

### Extended data figures

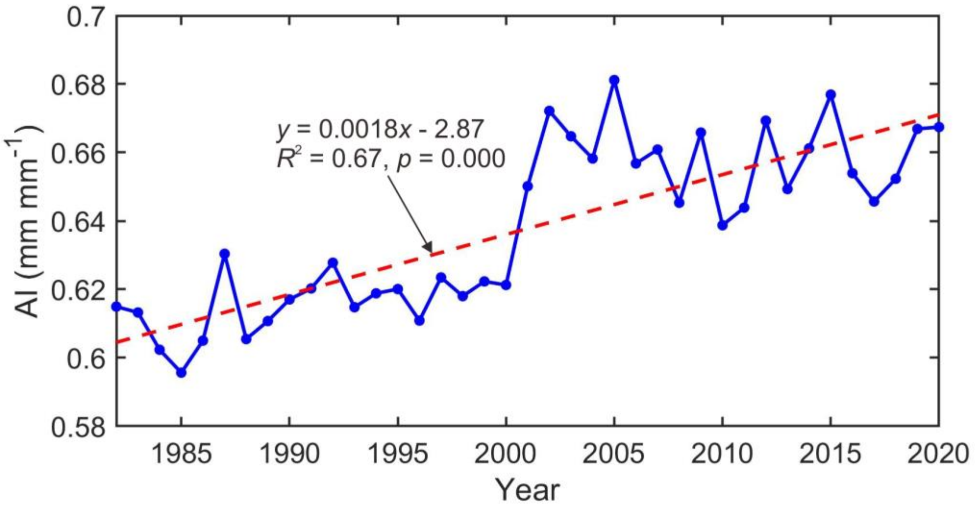


**Extended Data Fig. 1.** Change of arid index (AI) in 1982-2020.

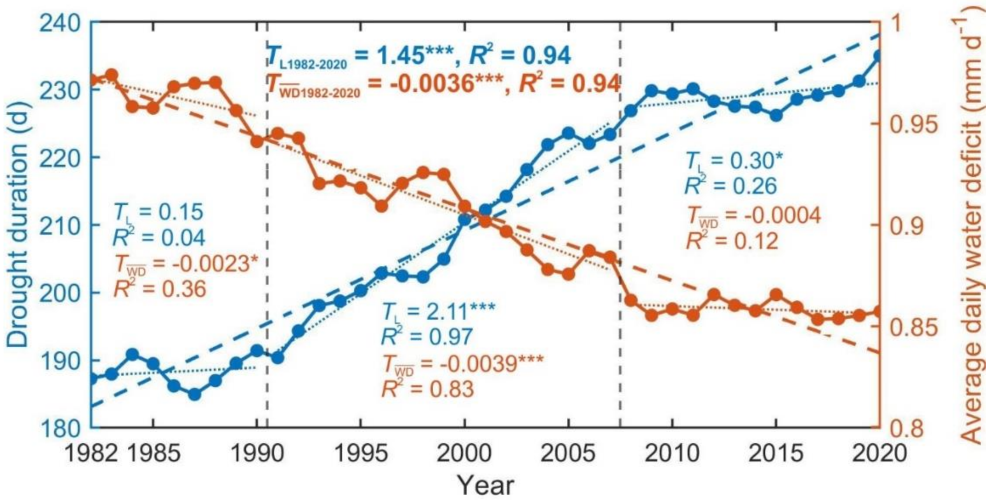


**Extended Data Fig. 2.** The trends of drought duration and average daily water deficit in global vegetated land from 1982 to 2020. The time period is divided into three segments: 1982-1990, 1991-2007, and 2008-2020, corresponding to the divisions in Fig.1. The trend lines $T_{\mathrm{L}}$ and $T_{\overline{\mathrm{WD}}}$ represent the changes in drought duration and average daily water deficit respectively. To indicate the significance of the regression coefficients, the following symbols are used: '*' indicating a significance level of 0.1, '**' indicating a significance level of 0.01, '***', and indicating a significance level of 0.001.

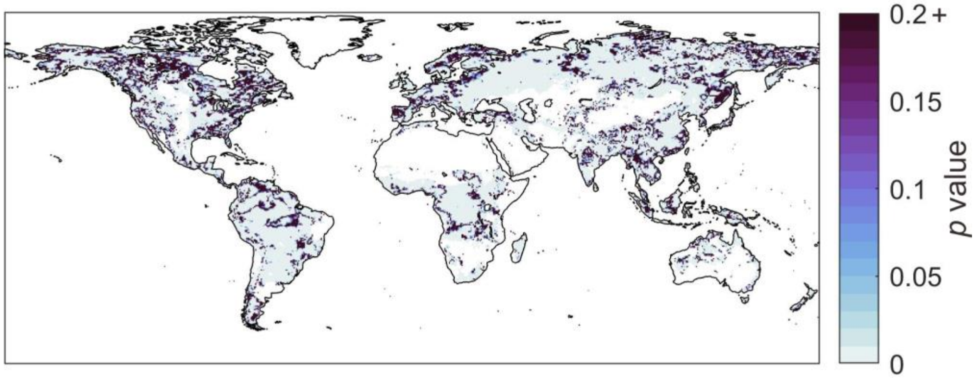


**Extended Data Fig. 3.** The *p* value distribution of $S_R$ changes with regression.

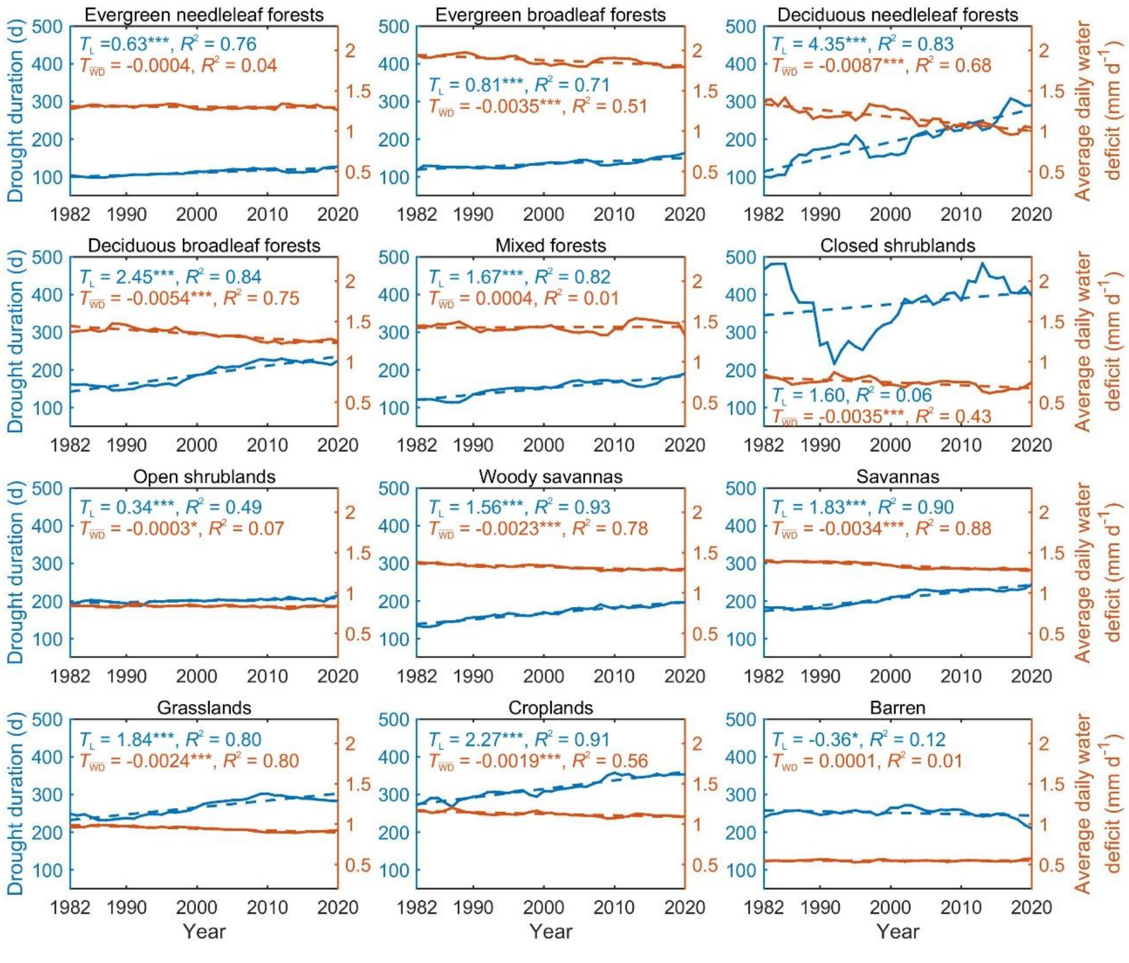


**Extended Data Fig. 4.** Trends of drought duration and average daily water deficit from 1982 to 2020 in different land cover types. The trend lines $T_L$ and $T_{\overline{WD}}$ represent the change of drought duration and average daily water deficit respectively. The significance of the regression coefficients uses the same symbols as in Extended Data Fig. 2.

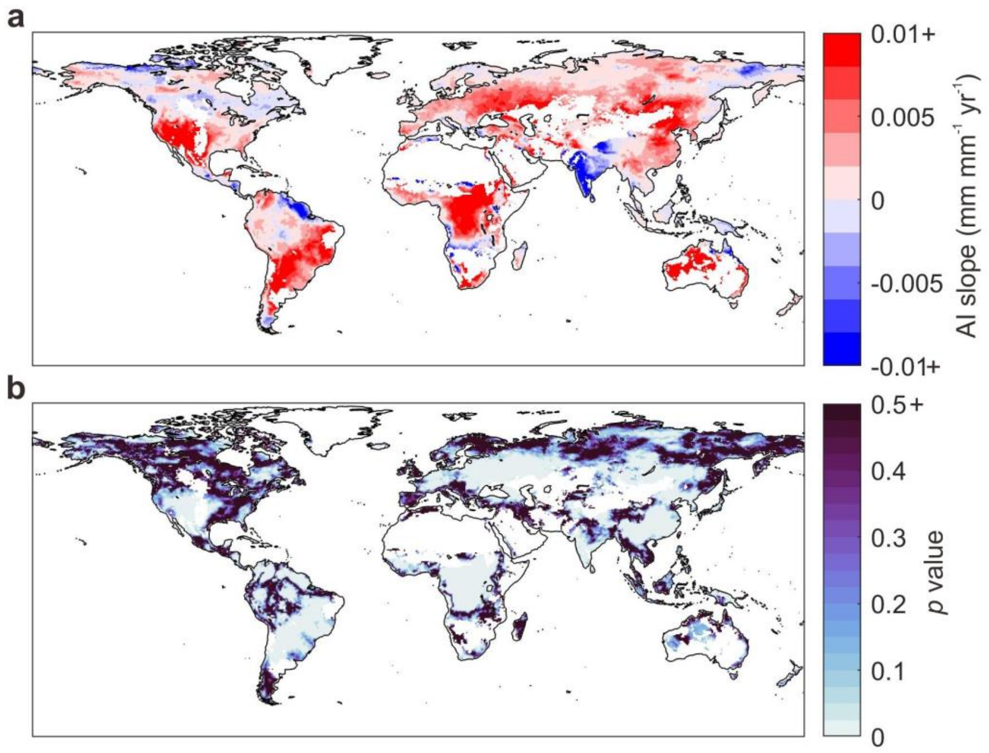


**Extended Data Fig. 5.** The spatial pattern of aridity index (AI) regression slope (**a**) and *p* value (**b**) in 1982-2020.

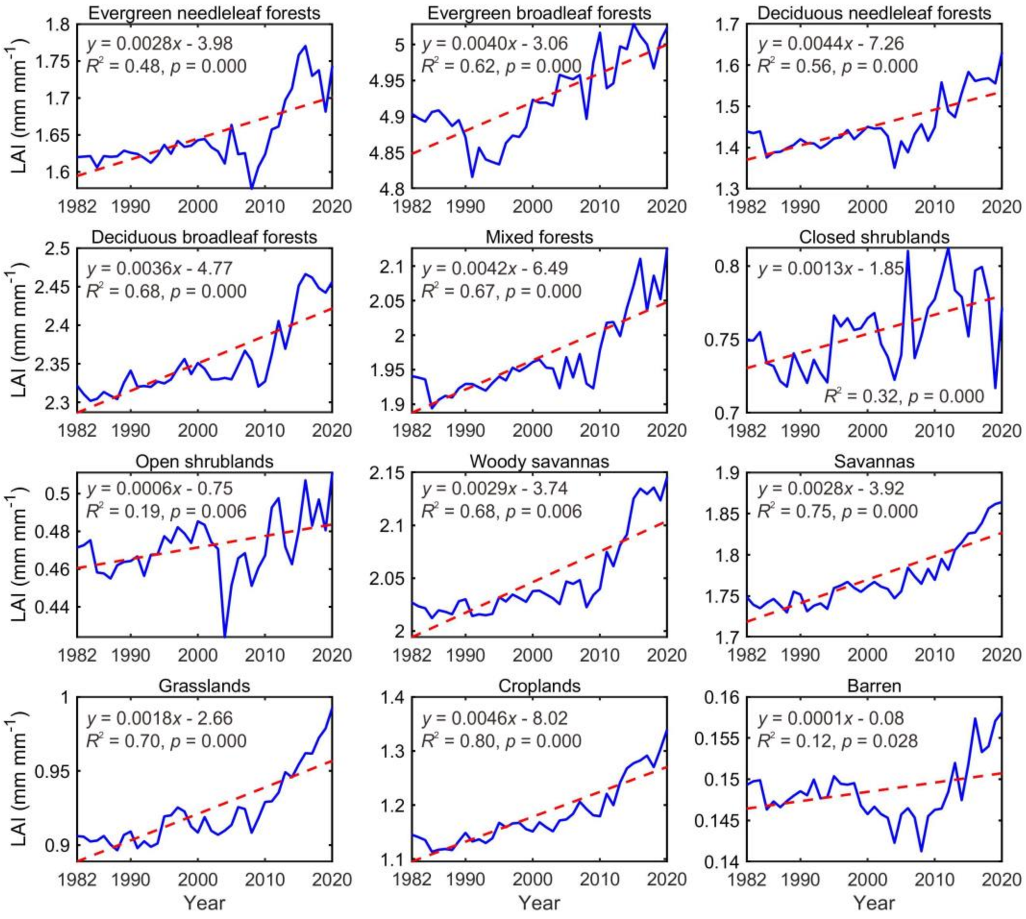


**Extended Data Fig. 6.** Change of leaf area index (LAI) in different land cover types in 1982-2020.

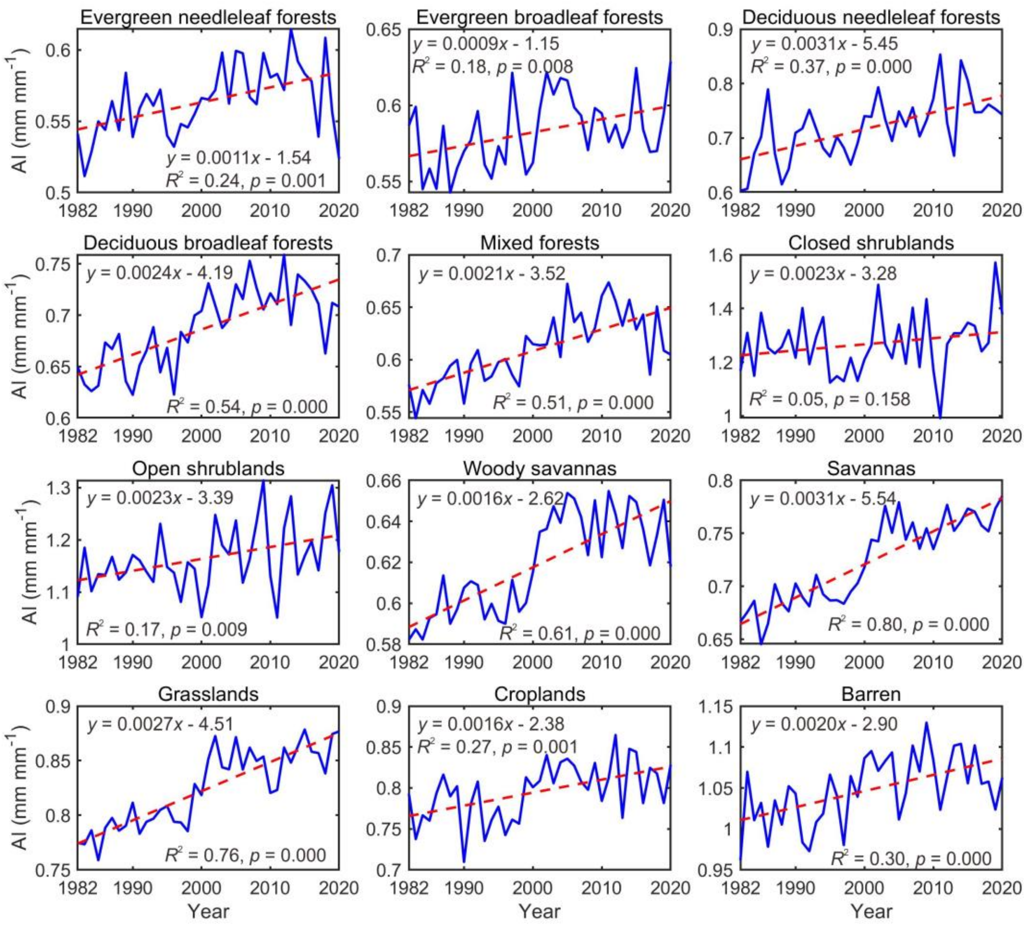


**Extended Data Fig. 7.** Change of arid index (AI) in different land cover types in 1982-2020.

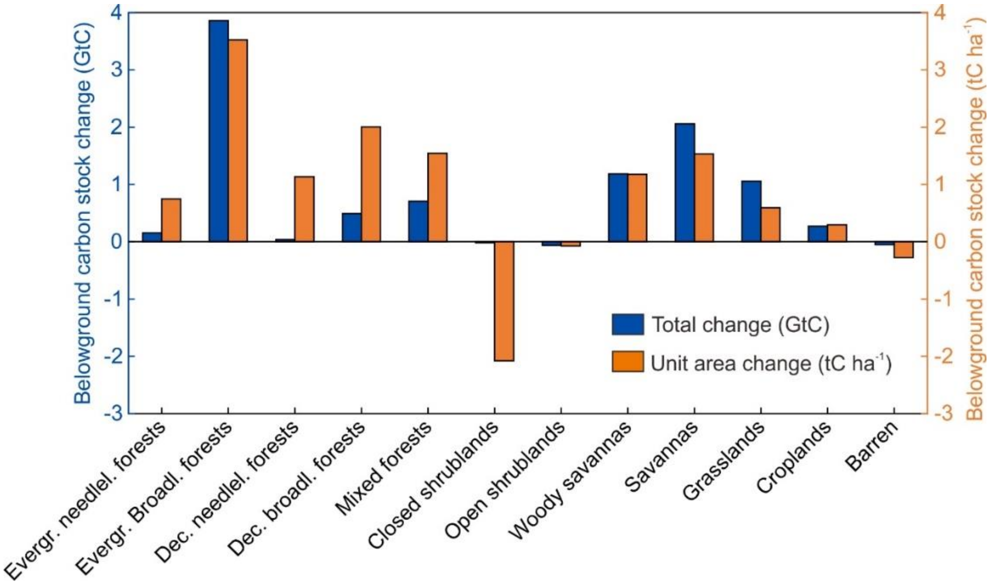


**Extended Data Fig. 8.** Change of belowground carbon stock in different land cover types in 1982-2020.

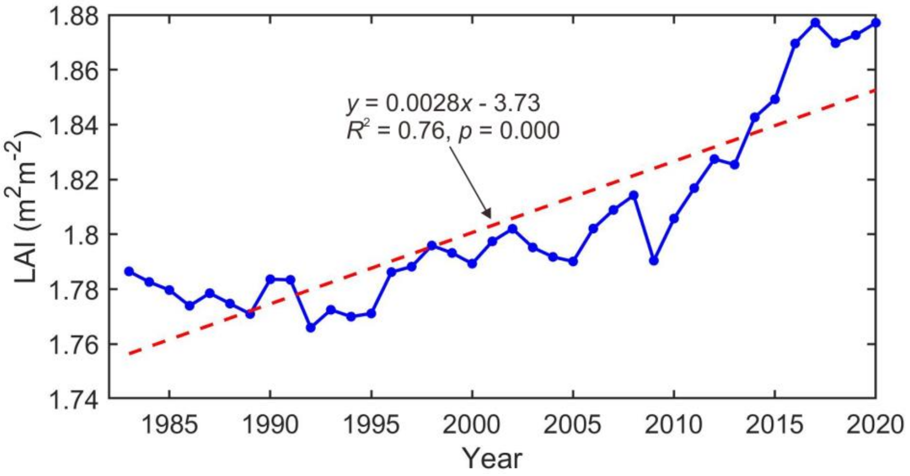


**Extended Data Fig. 9.** Change of global average LAI from 1982 to 2020.

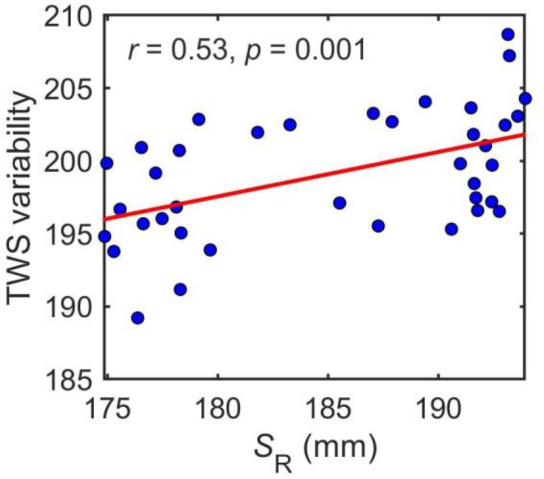


**Extended Data Fig. 10.** Spearman rank correlation between global average $S_R$ and annual TWS variability from 1982 to 2019.

**Supplementary information**

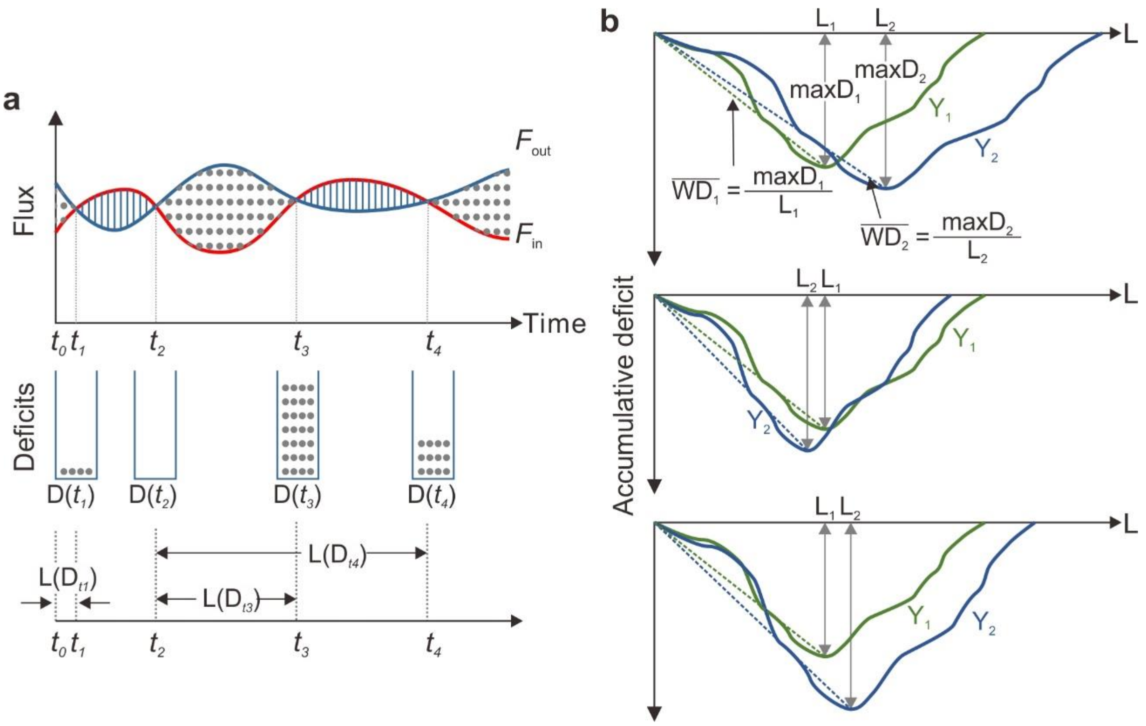


**Fig. S1.** Conceptual illustration of the algorithm for calculating $S_R$ (based on Wang-Erlandsson et. al., 2016) and analyzing the drivers of $S_R$ change:

(**a**) Algorithm for $S_R$ Calculation: The algorithm is based on the study by Wang-Erlandsson et al. (2016). It involves calculating the root zone storage capacity ($S_R$) by tracking the accumulative water deficit (*D*) over time. The shaded areas represent the differences (*A*) between outflow and inflow. *A* is positive when outflow is greater than inflow (dots shading) and negative when outflow is less than inflow (line shading). The accumulative water deficit *D* is increased by positive *A* and decreased by negative *A*. *D* is constrained to never become negative; when it reaches zero, it stays there. The accumulated deficit is illustrated by the number of dots in the containers. The accumulative period of *D* when it is greater than zero is termed drought duration (*L*).

(**b**) Algorithm for $S_R$ Change Attribution: In this study, $S_R$ is defined as the maximum value of *D* occurring during a period of 10 years, denoted as *maxD*, in mm. The accumulative deficit curve, ranges from zero to the maximum accumulation deficit (*maxD*) before returning to zero. The dashed line represents the average daily water deficit ($\overline{WD}$), in mm $d^{-1}$, during the drought duration *L*, in days. The illustration considers two critical periods in two different years, $Y_1$ and $Y_2$, representing year 1 and year 2, respectively. To analyze the drivers of $S_R$ change, the following comparisons are made: If $L_1<L_2$ and $|\overline{WD}_1|>|\overline{WD}_2|$, it indicates that the increase in $S_R$ is dominated by drought duration. If $L_1 > L_2$ and $|\overline{WD}_1|<|\overline{WD}_2|$, it suggests that the increase of $S_R$ is dominated by average daily water deficit. If $L_1< L_2$ and $|\overline{WD}_1|<|\overline{WD}_2|$, it implies that the increase of $S_R$ is influenced by both drought duration and average daily water deficit.

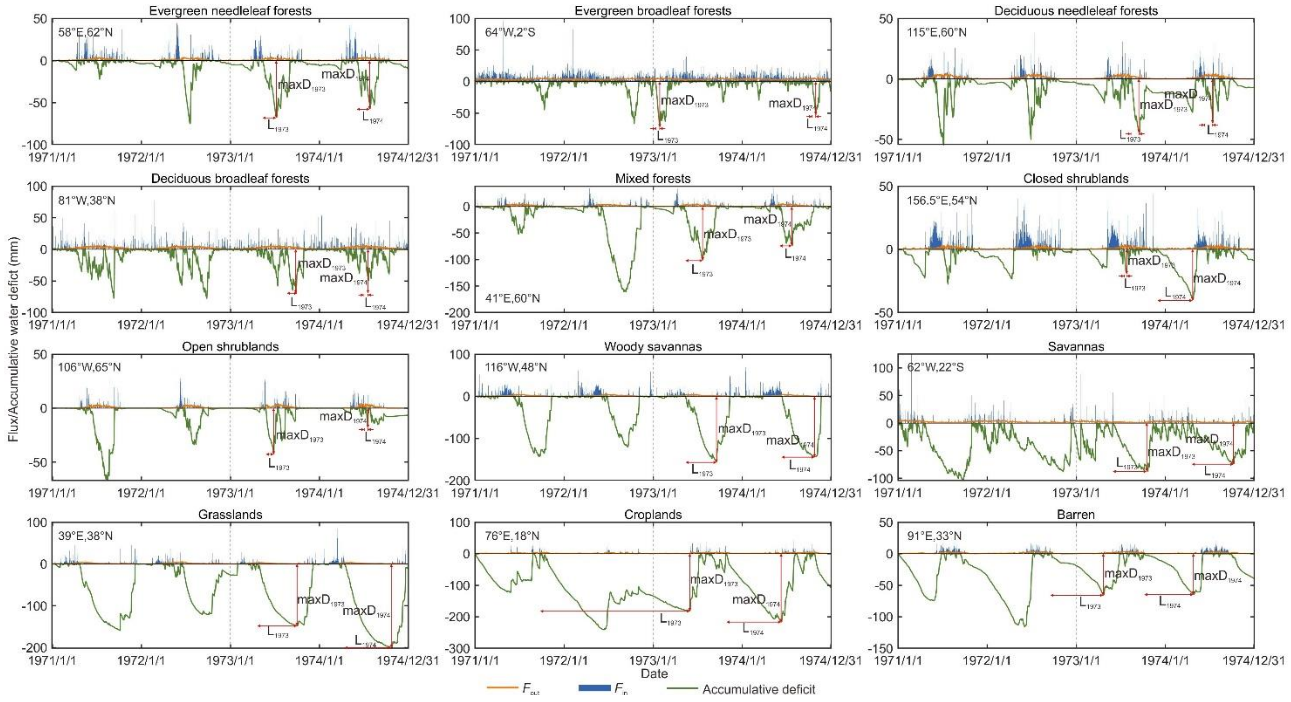


**Fig. S2.** Some examples of $S_R$ calculation at grids in different land cover types from 1971 to 1974. Outflow is evaporation, and inflow is the sum of precipitation, snowmelt and irrigation. *maxD* and *L* represents the maximum accumulative deficit and corresponding drought duration of the year. The subscript numbers represent the year 1973 and 1974, respectively. The data from 1971 to 1972 was used to warm-up (before the gray dashed line).

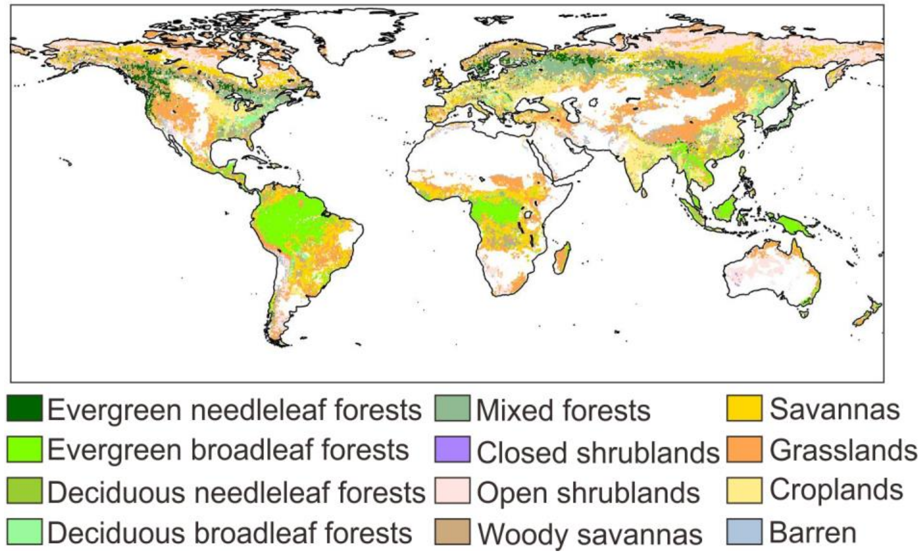


**Fig. S3** Distribution of land cover types.
